# Oxidation shuts down an auto-inhibitory mechanism of von Willebrand factor

**DOI:** 10.1101/786889

**Authors:** Rachel Tsai, Gianluca Interlandi

**Author notes:** Correspondence to: Gianluca Interlandi, Department of Bioengineering, University of Washington Box 355061, 3720 15^th^ Ave NE, Seattle, WA 98195-5061, USA.

## Abstract

The blood protein von Willebrand factor (VWF) is a key link between inflammation and pathological thrombus formation. In particular, oxidation of methionine residues in specific domains of VWF due to the release of oxidants in inflammatory conditions has been linked to an increased platelet-binding activity. However, the atomistic details how methionine oxidation activates VWF have not been elucidated to date. Yet understanding the activation mechanism of VWF under oxidizing conditions can lead to the development of novel therapeutics that target VWF selectively under inflammatory conditions in order to reduce its thrombotic activity while maintaining its haemostatic function. In this manuscript, we used a combination of a dynamic flow assay and molecular dynamics (MD) simulations to investigate how methionine oxidation removes an auto-inhibitory mechanism of VWF. Results from the dynamic flow assay revealed that oxidation does not directly activate the A1 domain, which is the domain in VWF that contains the binding site to the platelet surface receptor glycoprotein Ib*α* (GpIb*α*), but rather removes the inhibitory function of the neighboring A2 and A3 domains. Furthermore, the MD simulations combined with free energy perturbation calculations suggested that methionine oxidation may destabilize the binding interface between the A1 and A2 domains leading to unmasking of the GpIb*α*-binding site in the A1 domain.

## Introduction

The multimeric blood protein von Willebrand factor (VWF) is key in the initial stages of blood coagulation especially under high shear conditions.^1^ Once activated, VWF mediates the aggregation of platelets to the site of vascular injury.^1^ VWF is a relatively long multimeric chain, where each monomer consists of a number of domains. Under normal low shear conditions, VWF is thought to be coiled up and consequently does not significantly bind to platelets. However, anchorage to the exposed endothelium through its A3 domain or high shear cause VWF to unravel and expose the A1 domain, which contains the binding site to the platelet surface receptor glycoprotein Ib*α* (GpIb*α*).^2^ Tensile force generated by high shear also causes the A2 domain, located between A1 and A3, to unfold and expose a scissile bond that is cleaved by the metalloprotease ADAMTS13.^3^ This process converts VWF into smaller multimers that are less active in binding platelets. Recently, the platelet-binding function of VWF has been shown to be drastically enhanced in the presence of the oxidant hypochlorous acid (HOCl), which is produced under inflammatory conditions.^4^ The hyperactivity of VWF observed in the presence of HOCl has been linked to the conversion of methionine residues in the A1, A2 and A3 domains to methionine sulfoxide as observed in mass spectrometry experiments.^4^ Oxidation of methionine residues in VWF under oxidizing conditions has been reported also *in vivo* by a study involving patients with sickle cell disease,^5, 6^ a chronic inflammatory condition. These observations strongly suggest that conformational changes occur within VWF due to methionine oxidation, which contribute to a pro-thrombotic activity of VWF under inflammatory conditions.^7^

Because of its central role in thrombus formation, the protein VWF has been the target for anti-thrombotic interventions.^8, 9^ However, currently available anti-thrombotic therapies including the recently discovered caplacizumab, which directly inhibits VWF,^10^ are known to increase the risk of intracranial bleeding. Thus, there is a need to develop novel therapeutics that target VWF specifically under conditions that lead to thrombosis while maintaining its haemostatic function, i.e., the coagulation process that is needed to stop blood loss from a wound and promote healing. A major risk factor for pathological thrombus formation is inflammation.^11^ During the inflammatory response, neutrophils become activated^12^ and release hydrogen peroxide (H_2_O_2_), which is converted to HOCl through the action of myeloperoxidase (MPO). The oxidant HOCl is known to oxidize methionine residues in blood proteins altering their function.^13^ Understanding the structural mechanism behind the activation of VWF under oxidizing conditions resulting from inflammation can help foster the development of inhibitors of VWF that can differentiate between pathological thrombus formation and normal haemostasis.

There is strong evidence that VWF is regulated by an auto-inhibitory mechanism whereby binding of the A1 domain to GpIb*α* is inhibited by neighboring domains. Specifically, the A2A3 domains (located C-terminally to A1, Figure 1),^14, 15^ the D’D3 domains (located N-terminally, Figure 1),^16, 17^ and the linker between the D3 and A1 domain (called here “N-terminal linker”)^18–20^ have been found to decrease binding between A1 and GpIb*α*. Tensile forces generated by shear stress present in flowing blood are thought to remove the inhibitory function of neighboring domains of A1. It is likely that oxidation of methionine residues also reduces or eliminates the domain inhibition mechanism leading to the observed hyperactivation of VWF under oxidizing conditions.^4^ A previous molecular dynamics (MD) study by us suggested that methionine oxidation destabilizes the fold of the A2 domain such that it can no longer mask the GpIb*α*-binding site in the A1 domain.^21^ It is important to note that although under normal conditions unfolding of the A2 domain leads to its cleavage by ADATMS13 and downregulation of VWF, under oxidizing conditions the A2 domain cannot be cleaved because a methionine residue at position 1606 located in the scissile bond is converted to methionine sulfoxide.^22^ To date, no experimental evidence has been provided whether oxidizing conditions directly diminish the inhibitory function of the A2A3 domains on the ability of A1 to bind GpIb*α* since previous experiments have been performed with full-length VWF.^4, 22^

**Figure 1:**
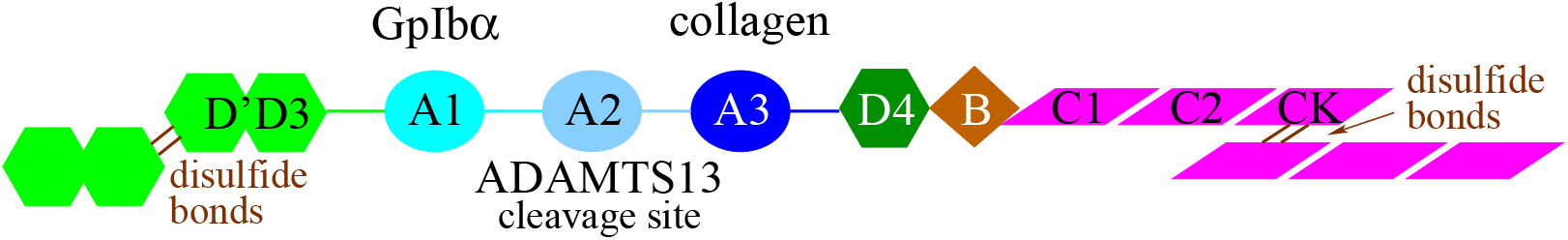
Overview of VWF structure. Schematic representation of a VWF monomer and function of the A1, A2 and A3 domains.

In this study, we used a dynamic flow-chamber based adhesion assay to evaluate whether oxidizing conditions reduce the inhibitory function of the A2A3 domains on A1. To this extent, we compared constructs consisting of the isolated A1 domain with constructs consisting of the A1A2A3 domains under oxidizing and non-oxidizing conditions. Furthermore, we used MD simulations to evaluate whether methionine residues located on the surface of the A1 and A2 domain destabilize the A1-A2 interface upon oxidation. Currently, no experimentally determined structure is available of any complex between A1 and a neighboring domain. However, computational docking studies of the A1A2 domains have been performed by others^23^ and the obtained models were used here as initial conformations for the MD simulations.

## Materials and Methods

### Expression of recombinant VWF constructs

The DNA constructs for isolated VWF A1 domain (residues 1238-1472) and VWF A1A2A3 (1238-1878) were obtained as previously described for the isolated A1 domain.^18^ To facilitate specific anchoring on a surface, the constructs contained at the C-terminus an additional four repeats of GGGGS linker followed by a biotin tag with amino acids LNDIFEAQKIEWH. The recombinant expression constructs were transfected into CHO Tet-On cells and expression was carried out in the presence of biotin as previously described.^18^ Culture medium containing secreted recombinant A1 or A1A2A3 protein was collected and concentrated five times using an Amicon Ultra-15 Centrifugal Filters (EMD Millipore, Billerica, MA). Free biotin was removed from the samples using a desalting PD-10 column (GE Healthcare, Marlborough, MA).

### Preparation of surfaces with VWF constructs

VWF constructs were anchored to surfaces via the biotin tag similarly to a previous study by us.^18^ Briefly, 35-mm tissue culture polystyrene dishes were incubated with a 100 *μ*l droplet of 200 *μ*g/ml biotin-BSA for 1.5 hours at 37° C, washed 3 times with 0.2% BSA-PBS (w/v), incubated with 100 *μ*g/ml streptavidin for 30 minutes at 37° C, washed 3 times with 0.2% BSA-PBS (w/v) and incubated over night at 4° C in 0.2% BSA-PBS (w/v). Dishes were then incubated for 30 minutes at 37° C with a 100 *μ*l solution containing the biotinylated VWF A1 or A1A2A3 prepared as described above and washed 3 times with 0.2% BSA-PBS (w/v). The incubation with A1 or A1A2A3 was repeated two more times for a total of three incubations. The plates were then blocked with 0.2% BSA-PBS (w/v). The amount of biotinylated VWF used in this surface functionalization was determined to be sufficient to saturate the available biotin-binding sites by using biotin-4-fluorescin to test for the presence of free biotin-binding sites using the method of Kada et al^24^ as detailed in a previous study by us.^18^

### Oxidation of VWF constructs

Oxidation of the VWF constructs was achieved by incubating the plates (with the constructs anchored as described above) with 5 *μ*M HOCl for 30 minutes at 37° C. We estimated that this concentration yields a molar ratio of 1:50 of VWF protein molecules to oxidant, which is of similar magnitude as the molar ratios present in a previous oxidation study.^4^ The oxidation reaction was quenched by adding free methionine.

### Preparation of GPIb*α*-coated beads

The N-terminal domain of GPIb*α* (amino acids 1-300 of the extracellular domain of GPIb*α* with a C-terminal biotin tag for oriented anchoring) was made using a previously described expression vector^25^ and expressed in CHO cells. After desalting and concentrating via ammonium sulfate precipitation, the GC300 was bound to 3 *μ*m-diameter streptavidin-coated Dynabeads (Invitrogen) with gentle mixing for 30 minutes at room temperature. Beads were then washed and resuspended in phosphate-buffered saline (PBS) with 0.2% BSA (w/v) for use in the flow chamber.

### Beads adhesion in flow

Platelet adhesion studies were performed in a GlycotechTM parallel plate flow chamber as previously described,^18^ using functionalized 35-mm dishes prepared as described above as the lower surface. A 300-*μ*l bolus of GPIb*α*-coated beads was introduced into a flow chamber and allowed to settle for 30 seconds, before 0.2% BSA-PBS (w/v) buffer was pushed through the chamber at the indicated shear stress and beads observed with a 10X objective and CCD camera. Time-lapse videos were taken with a 10X objective using the program ImageJ, at 1 frame per second, and the beads in the videos tracked using SVCell RS by DRVision Technologies LLC, Bellevue WA. The tracks were processed using a MATLAB script to determine the number of beads bound at each recorded frame as previously described.^18^ For each shear stress, the beads were washed over the surface for 30 seconds and the number of bound beads was averaged over this time window. The fraction of bound beads was then calculated by dividing the average number of bound beads at a given shear stress by the number of beads initially bound. The latter was determined by averaging the first 10 seconds of the 30-seconds time window where the beads were washed at 4 dyne/cm^2^.

### Initial conformations for MD simulations

The initial conformations for the simulations with isolated A1 and A2 domain were derived from the respective crystallographic structures: PDB code 1AUQ for A1 and PDB code 3GBX (chain A) for A2, respectively. Some of the simulations with the A2 domain were already performed as part of previous studies^21, 26^ as noted in Table 1. The simulations with the A1A2 domains complex were started from six models, which were obtained in a previous study by others^23^ using a combination of patchdock^27^ and firedock.^28, 29^ The coordinates of the models were kindly provided to us by the authors. For the purposes of this manuscript, the models are labeled, A1A2 n, where n is a number between 1 and 6. All initial conformations were minimized with 100 steps of steepest descent in vacuo using the program CHARMM.^30^

**Table 1.** Simulation Systems.

| **Name** | **Starting structure** | **Type** | **Duration [ns]** |
| --- | --- | --- | --- |
| A1_sim_1,2 | PDB code 1AUQ | 300 K<sup>a</sup> | 2 x 50 |
| A2_sim_1,2,3 | PDB code 3GXB | 300 K<sup>a</sup> | 3 x 40<sup>b</sup> |
| A1A2_1_sim1,2 | complex model | 300 K<sup>a</sup> | 2 x 50 |
| A1A2_2_sim1,2 | complex model | 300 K<sup>a</sup> | 2 x 50 |
| A1A2_3_sim1,2 | complex model | 300 K<sup>a</sup> | 2 x 50 |
| A1A2_4_sim1,2 | complex model | 300 K<sup>a</sup> | 2 x 50 |
| A1A2_6_sim1,2 | complex model | 300 K<sup>a</sup> | 2 x 50 |
| A1A2_6_sim1,2 | complex model | 300 K<sup>a</sup> | 2 x 50 |
| A1_M1393_fep_1,2,3 | A1_WT_1 (10+20 ns), A1_WT_2 (10 ns) | FEP | 3 x 10 |
| A2_M1495_fep_1,2,3 | A2_WT_1 (10+20 ns), A2_WT_2 (10 ns) | FEP | 3 x 10<sup>c</sup> |
| A2_M1545_fep_1,2,3 | A2_WT_1 (10+20 ns), A2_WT_2 (10 ns) | FEP | 3 x 10 |
| A1A2_1_M1495_fep_1,2,3 | A1A2_2met_1 (10+20 ns), A1A2_2met_2 (10 ns) | FEP | 3 x 10 |
| A1A2_2_M1393_fep_1,2,3 | A1A2_2met_1 (10+20 ns), A1A2_2met_2 (10 ns) | FEP | 3 x 10 |
| A1A2_2_M1495_fep_1,2,3 | A1A2_2met_1 (10+20 ns), A1A2_2met_2 (10 ns) | FEP | 3 x 10 |
| A1A2_3_M1545_fep_1,2,3 | A1A2_1met_1 (10+20 ns), A1A2_1met_2 (10 ns) | FEP | 3 x 10 |
| A1A2_4_M1393_fep_1,2,3 | A1A2_2met_1 (10+20 ns), A1A2_2met_2 (10 ns) | FEP | 3 x 10 |
| A1A2_4_M1495_fep_1,2,3 | A1A2_2met_1 (10+20 ns), A1A2_2met_2 (10 ns) | FEP | 3 x 10 |
| A1A2_5_M1545_fep_1,2,3 | A1A2_1met_1 (10+20 ns), A1A2_1met_2 (10 ns) | FEP | 3 x 10 |
| A1A2_6_M1545_fep_1,2,3 | A1A2_1met_1 (10+20 ns), A1A2_1met_2 (10 ns) | FEP | 3 x 10 |

<sup>a</sup>These simulations sampled the unoxidized conformational state of either the isolated domains or the A1A2 domains complex at room temperature and did not involve FEP. <sup>b</sup>These simulations were performed as part of a previous study.<sup>26</sup> <sup>c</sup>These simulations were performed as part of a previous study.<sup>21</sup> All simulations were performed in either duplicates or triplicates and labeled with indexes (e.g., 1, 2, 3 for triplicates), respectively. The FEP simulations were started from snapshots sampled along the "300 K" runs as indicated.

### General setup of the systems

The MD simulations were performed with the program NAMD^31^ using the CHARMM all-hydrogen force field (PARAM22)^32^ with the CMAP extension^33, 34^ and the TIP3P model of water. The force field parameters for methionine sulfoxide optimized for the PARAM22 force field were downloaded from the SwissSidechain website.^35^ The different simulation systems are summarized in Table 1. The proteins were inserted into a cubic water box with a side length of 84 Å for the A1 domain and 100 Å for the A1A2 complex, respectively, resulting in systems with in total ca. 56,000 and ca. 95,000 atoms, respectively. Most of the simulations with the A2 domain discussed here were performed as part of previous studies^21, 26^ and were re-analysed for the purposes of this manuscript (see Table 1 for detail). The water molecules overlapping with the protein were removed if the distance between any water atom and any atom of the protein was smaller than 2.4 Å. Chloride and sodium ions were added to neutralize the system and approximate a salt concentration of 150 mM. To avoid finite size effects, periodic boundary conditions were applied. After solvation, the system underwent 500 steps of minimization while the coordinates of the heavy atoms of the protein were held fixed and subsequent 500 steps with no restraints. Each simulation was started with different initial random velocities to ensure that different trajectories were sampled whenever the same starting conformation was used. Electrostatic interactions were calculated within a cutoff of 10 Å, while long-range electrostatic effects were taken into account by the Particle Mesh Ewald summation method.^36^ Van der Waals interactions were treated with the use of a switch function starting at 8 Å and turning off at 10 Å. The dynamics were integrated with a time step of 2 fs. The covalent bonds involving hydrogens were rigidly constrained by means of the SHAKE algorithm with a tolerance of 10^−8^. Snapshots were saved every 10 ps for trajectory analysis.

### Equilibration and sampling of the native state

Before production runs, harmonic constraints were applied to the positions of all heavy atoms of the protein to equilibrate the system at 300 K during a time length of 0.2 ns. After this equilibration phase, the harmonic constraints were released and simulation runs were performed for a total of 50 ns each (Table 1). The first 10 ns of unconstrained simulation time were also considered part of the equilibration and were thus not used for the analysis. During the equilibration and in all production runs, the temperature was kept constant at 300 K by using the Langevin thermostat^37^ with a damping coefficient of 1 ps^−1^, while the pressure was held constant at 1 atm by applying a pressure piston.^38^

### Change in free energy of binding upon oxidation

The change in the free energy of binding between the A1 and A2 domain due to the oxidation of a methionine residue was estimated by making use of alchemical transformations^39^ in combination with the thermodynamic cycle^40^ represented in Figure 2. The alchemical transformations were performed through FEP calculations.^41^ Each alchemical transformation was performed in the forward and backward direction. In the forward transformation, a methionine side chain is slowly converted to methionine sulfoxide, which contains an oxygen atom covalently bound to the sulphur atom. The conformation achieved in the forward transformation is then used to start a backward transformation where methionine sulfoxide is converted back to methionine. During the process, the amount of work needed for each transformation is calculated. The forward and backward calculations were then combined and a value for the ΔG of the oxidation reaction was obtained using the Bennett’s acceptance ratio method^42^ implemented in the ParseFEP plugin of VMD.^43^ Each forward and backward transformation was performed for 10 ns (20 ns in total) during which a parameter *λ* was varied from 0 (unoxidized methionine) to 1 (methionine sulfoxide) and from 1 to 0, respectively, in time intervals of the length of 0.1 ns for a total of 100 intermediate states for each direction. The first half of each time window involved equilibration and the second half data collection. A soft core term was introduced to avoid singularities in the van der Waals potential.^44^ Each transformation (forward and backward) was performed in triplicates to evaluate statistical significance. In a previous study with the isolated A2 domain, similar ΔG values were obtained when the alchemical transformation was performed for only 1 ns and with time intervals of 0.025 ns^21^ indicating that the side chain has likely thoroughly sampled its local environment in the nanosecond time scale. Alchemical transformations were performed in either the isolated A1 or A2 domain, respectively (depending in which domain the side chain is located), and in the A1A2 domains complex. The calculations were performed in triplicates using snapshots sampled along the 50-ns (40-ns in the case of A2) simulations as initial conformations (Table 1), and the results were averaged. By considering the thermodynamic cycle (Figure 2), the difference in the free energy of binding upon methionine oxidation can be approximated by the difference between the ΔG values calculated from the alchemical transformations in the isolated (unbound) and complex (bound) state (see caption of Figure 2).

**Figure 2:**
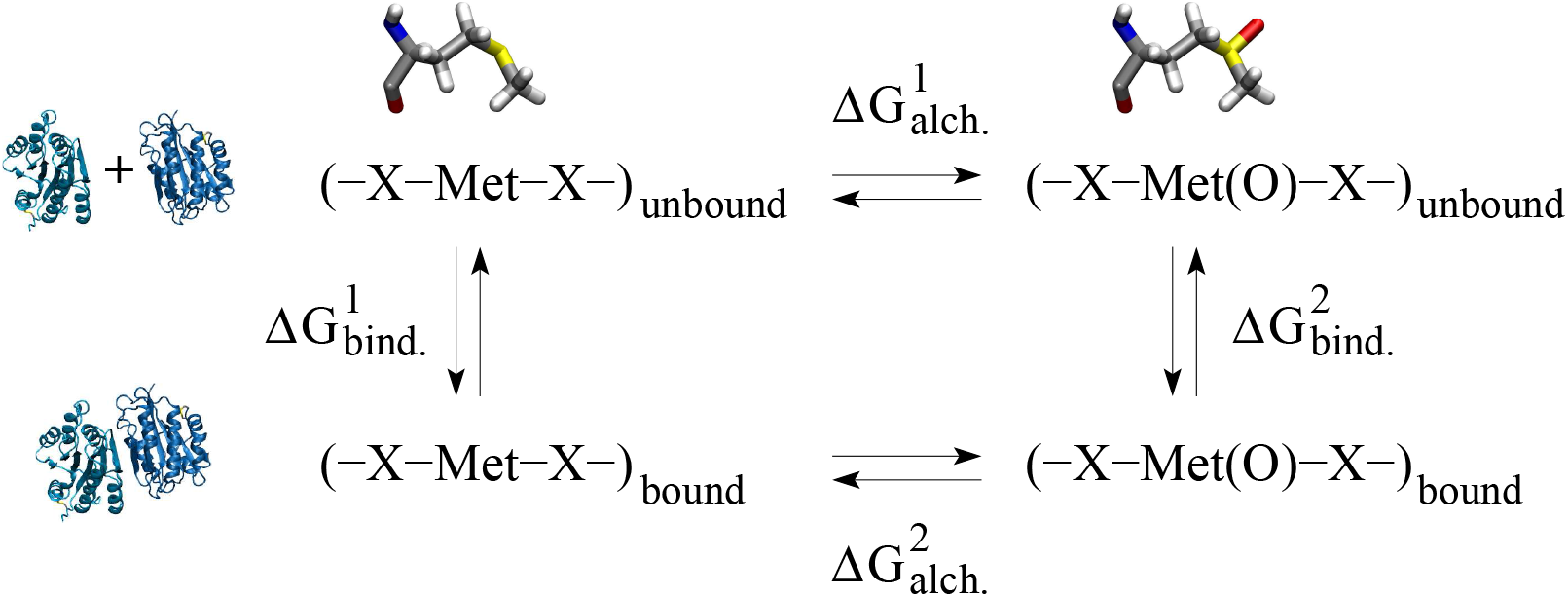
Thermodynamic cycle used to estimate the change in free energy of binding due to the oxidation of a methionine residue. The horizontal arrows correspond to the alchemical transformation from methionine to methionine sulfoxide (illustrated at the top) in the unbound (isolated A1 or A2 domain) and bound (A1A2 complex) state, respectively (illustrated on the left side). The vertical arrows describe the binding process in the unoxidized and oxidized state, respectively. 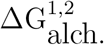 is calculated as described in the text. The change in free energy of binding can then be derived as follows: 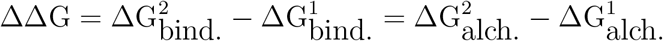.

### Determining methionine residues located at the interdomain interface and surface buried between the two domains

To determine whether a methionine residue was located at the inter-domain interface of a particular A1A2 complex model, we used the coordinates of the initial conformation for that model. If any side chain atom of a particular methionine of one domain was within 4 Å of any atom of a side chain located on the other domain, then that methionine was deemed to be at the interdomain interface. According to this criterion, M1393 was located at the interdomain interface of A1A2 2 and A1A2 4, M1495 at the interface of A1A2 1, A1A2 2 and A1A2 4, and M1545 at the interface of A1A2 3, A1A2 5 and A1A2 6, respectively (Figure 3).

**Figure 3:**
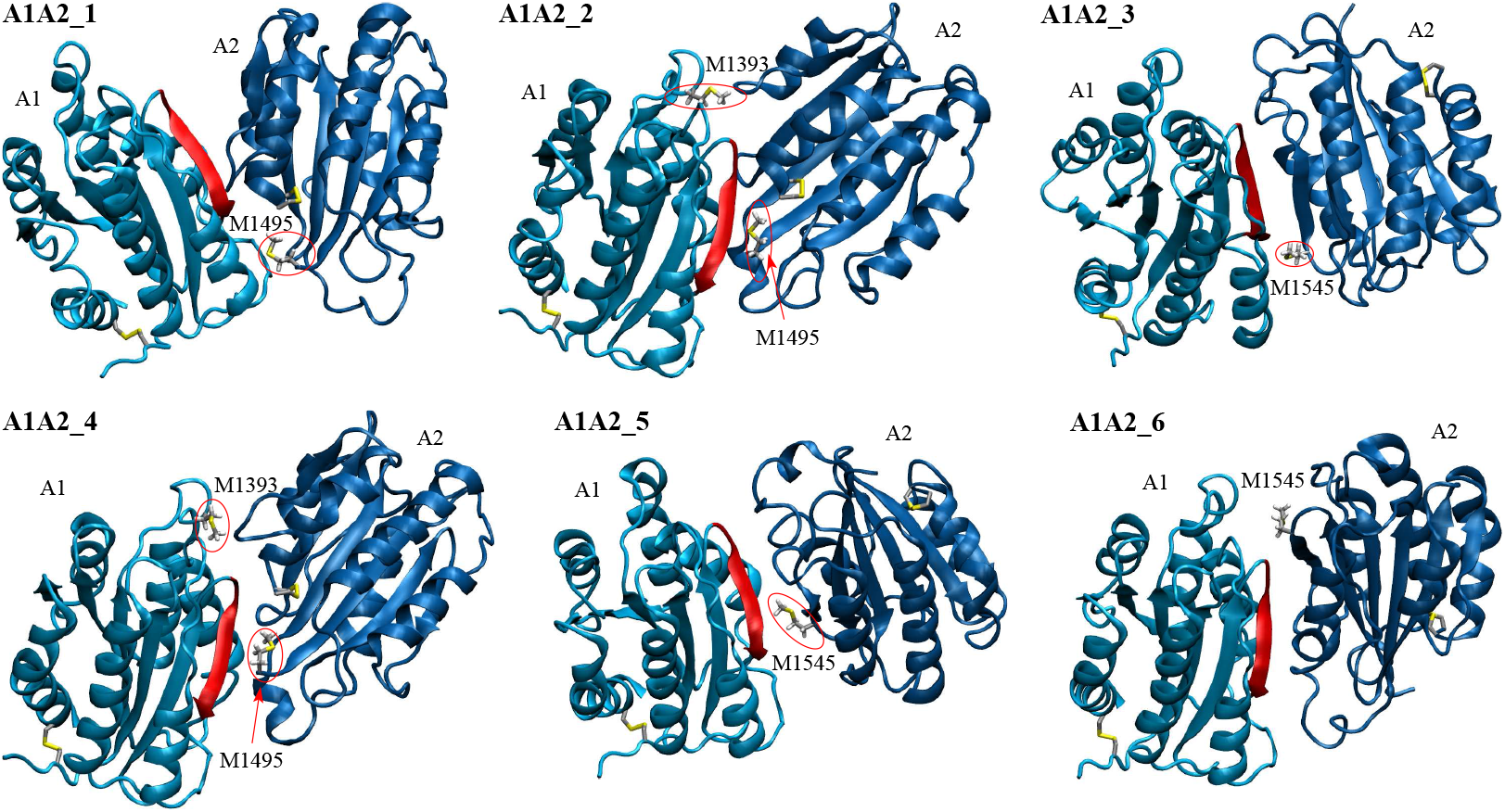
Models of the A1A2 domains complex. Methionine residues located near the interface and cysteine side chains forming disulfide bonds are shown in the stick and ball representation. Methionine residues at the interface are also labeled and highlighted by red circles. The backbone in the A1 domain colored in red highlights one of the major contact sites to GpIb*α*.^54^

The SASA buried at the inter-domain interface in a A1A2 domains complex along the MD trajectories was calculated using the program CHARMM^30^ by adding the SASA of each domain isolated and subtracting the SASA of the entire complex.

## Results

### Flow chamber experiments

We first investigated whether oxidizing conditions directly enhance binding between the isolated A1 domain and GpIb*α*, or whether oxidation removes the inhibitory function of the neighboring A2A3 domains. For this reason, the following two constructs were expressed: the isolated A1 domain and A1A2A3. Each construct was anchored to a surface through its C-terminal linker while the binding domain of GpIb*α* was anchored to beads as described in the Methods section. After anchoring them to the surface, constructs were either left in their normal unoxidized state, or were subjected to oxidizing conditions as described in the Methods section. The beads were allowed to settle onto the surface, washed at 2 dyne/cm^2^ fluidic wall shear stress, and then subjected to stepwise increasing or decreasing flow while their movement was recorded with videomicroscopy.

Oxidation of the isolated A1 domain did not alter the fraction of bound beads at any shear (Figure 4a). However, the oxidized A1A2A3 construct caused at high shear a larger fraction of beads to remain bound, than the non-oxidized construct (Figure 4b,c). Specifically, at high shear, we observed an up to 68% increase in the fraction of bound beads upon oxidation of A1A2A3 (Figure 4c). These results indicate that oxidation is likely to reduce the inhibitory effect that the A2A3 domains have on the A1 domain.

**Figure 4:**
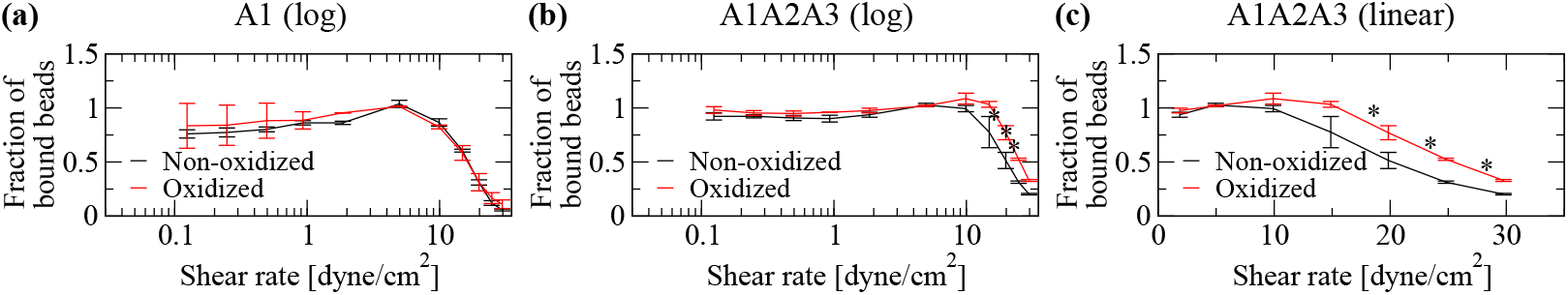
Flow chamber experiments. Fraction of bound beads to beads initially bound (see “Materials and Methods”). (a) Isolated A1 domain (logarithmic scale). (b) Construct containing VWF domains A1, A2 and A3 (logarithmic scale). (c) Same as in (b) but in linear scale to emphasize differences at intermediate and high shear (values below 1 dyne/cm^2^ were omitted to simplify graph appearance). Oxidation was performed using 5 *μ*M HOCl. Reported are averages and standard errors of the mean for two measurements. Statistical analysis was performed using a Student’s t-test. A difference with a p-value smaller than 0.05 was considered to be statistically significant and is indicated by an asterisk (*).

### Change in free energy of binding between A1 and A2 due to oxidation

The experiments performed here highlight that oxidation removes an auto-inhibitory mechanism of VWF, which consists of the A2A3 domains inhibiting binding of A1 to GpIb*α*. Understanding the atomistic mechanism behind the oxidation-induced activation of VWF can foster the development of therapeutics that target VWF in particular under inflammatory conditions, which are known to increase the risk of thrombosis.^11^ Mass spectrometry studies have shown that the presence of HOCl facilitates the conversion of methionine residues in the A1, A2 and A3 domains to methionine sulfoxide^5^ in particular in the presence of shear stress.^4^ Furthermore, VWF in blood samples from patients suffering of sickle cell disease, an inflammatory condition, has been found to have a higher rate of methionine oxidation compared to VWF from healthy donors.^5, 6^ This indicates that methionine oxidation in VWF due to oxidizing conditions is likely to play a physiological role. A previous MD study by us suggested that oxidation of methionine residues buried within the A2 domain decreases its thermodynamic stability favoring the unfolded state.^21^ However, it is also likely that oxidation of methionine residues located at the interface between the A1 and A2 domains plays a role in removing the inhibitory function that A2 has on the A1 domain. This hypothesis was explored here with MD simulations of the A1A2 domains complex and free energy perturbation calculations.

Currently, no experimental structure is available of the A1A2 domains complex. However, a previous computational study provided six models for the A1A2 domains complex.^23^ Analysis of the modeled structures revealed that in all six models at least one methionine residue of either the A1 or A2 domain, or both was located at the binding interface (see “Materials and Methods” and Figure 3). Furthermore, in all models the domains were oriented in such a way that the A2 domain obstructs the GpIb*α* binding site in the A1 domain,^23^ which is consistent with experimental studies showing that the A2 domain inhibits or reduces binding of A1 to blood platelets.^14, 15^ In order to study the stability of the interface of the A1A2 complex and generate initial conformations for free energy perturbation calculations, two 50-ns long MD simulations were performed at 300 K for each model of the complex (Table 1). The total aggregated simulation time for the A1A2 complex including FEP was 1.08 *μs*.

The six complex models were generally stable with the C_*α*_ root mean square deviation (RMSD) from the initial conformation on average below Å in most simulations (Figure 5a and Supplementary Figure 1). The exception was one simulation with A1A2 5 and one simulation with A1A2 6 where the two domains slightly unhinged away from each other although inter-domain contact was still maintained as indicated by the SASA buried at the interface (Figure 5b and Supplementary Figure 2). In order to estimate the difference in free energy of binding between the A1 and A2 domain due to oxidation of methionine residues located at the inter-domain interface, three snapshots were taken along the trajectories with each of the A1A2 models. Then, FEP calculations were performed (Table 1) where each methionine residue deemed to be located at the interface (see “Materials and Methods” and Figure 3) was individually transformed into methionine sulfoxide (see “Materials and Methods”). The transformations were also performed separately in the isolated A1 and A2 domain (Table 1) in order to calculate the difference in free energy of binding (ΔΔG) due to the oxidation of a methionine residue according to the thermodynamic cycle (Figure 2). The results revealed that oxidation of M1545 in A1A2 3, M1495 in A1A2 4 and M1545 in A1A2 6 had a statistically significant effect while oxidation of M1495 in A1A2 1 and M1393 in A1A2 2 had a marginally statistically significant effect in reducing the free energy of binding between A1 and A2 (Figure 6a,b). Overall, methionine oxidation had either a statistically or marginally statistically significant effect in five out of the six complex models in destabilizing the interface (Figure 6a,b). The destabilizing effect was strongest for M1545 in A1A2 3 probably due to the methionine residue being almost entirely buried at the interface (Figure 7). In fact, the sulfinyl group in methionine sulfoxide renders the side chain more hydrophilic than methionine, and thus its burial is energetically less favorable due to the loss of interaction with water molecules. In summary, these results suggest that methionine oxidation not only destabilizes the fold of the A2 domain as indicated by a previous study,^21^ but it is likely to also directly impair binding between the A1 and A2 domain.

**Figure 5:**
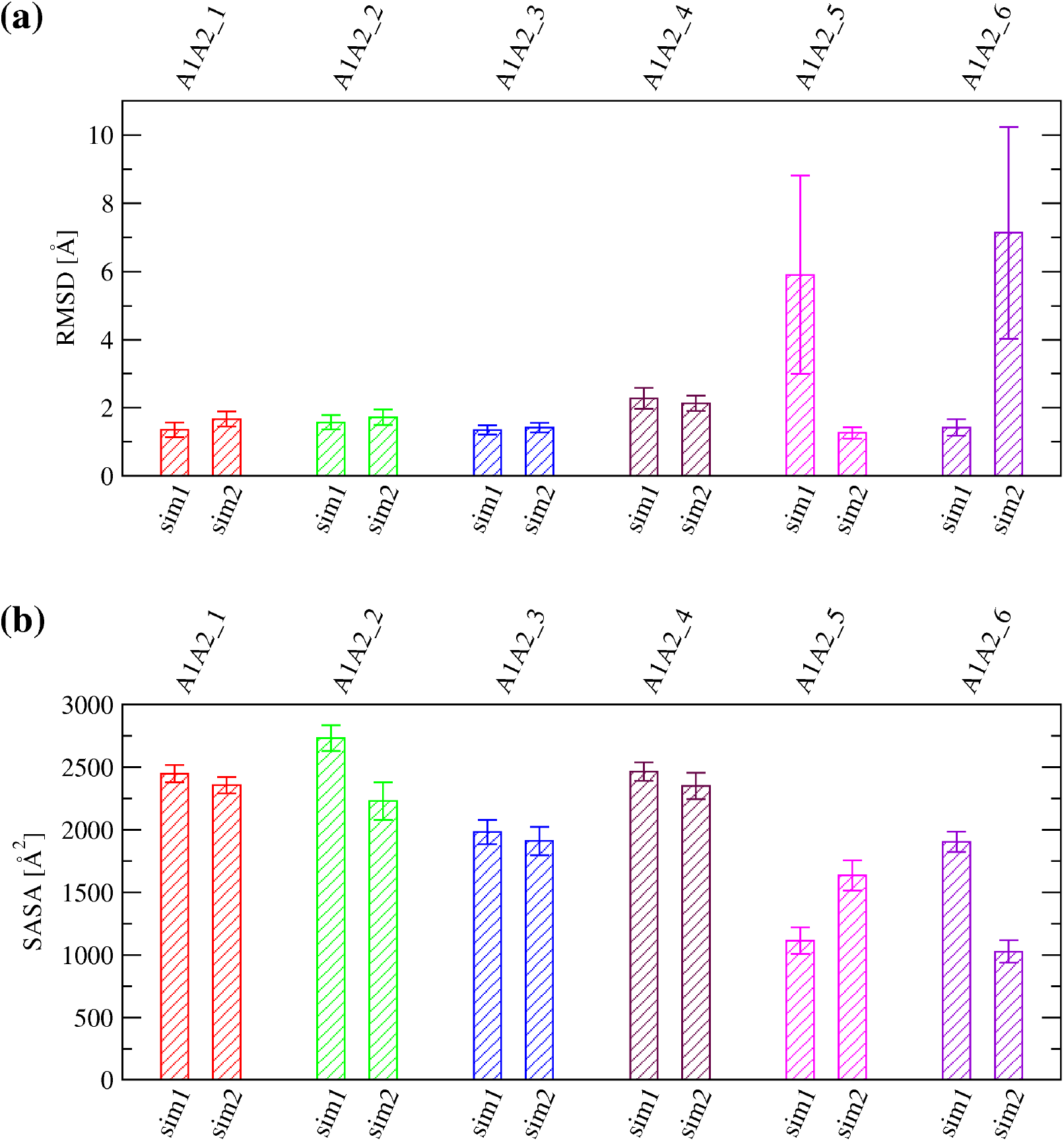
Stability of the A1A2 complex along room temperature simulations. (a) Averages and standard deviations of the C_*α*_ RMSD from the initial conformation calculated over the last 40 ns of in total 50-ns long simulations performed at 300 K with the A1A2 complex models. (b) Averages and standard deviations of the SASA buried at the inter-domain interface between A1 and A2 calculated over the last 10 ns of in total 50-ns long simulations with the complex models. The labels “sim1” and “sim2” refer to the two independent simulations run for each A1A2 complex model.

**Figure 6:**
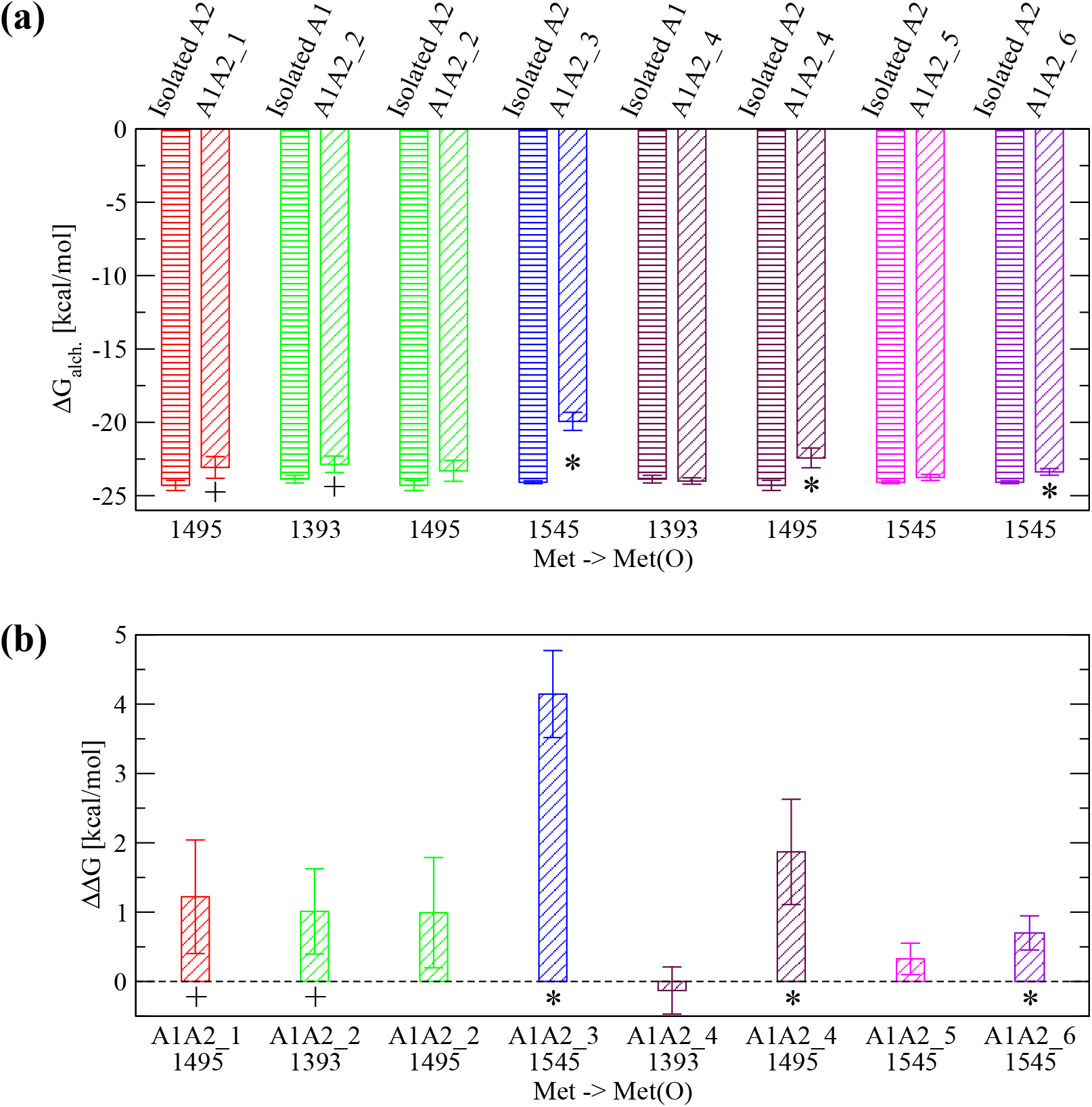
Estimate for the change in free energy of binding upon methionine oxidation. (a) Calculated ΔG_alch._^1,2^ for the transformation of a given methionine residue to methionine sulfoxide using FEP (see Figure 2 for details). Each transformation was performed in either the isolated A1 or A2 domain, respectively (bars with horizontal lines) and in the A1A2 complex model where the specific methionine residue is located at the interface (bars with oblique lines). The protein structure (isolated A1 or A2, or A1A2 complex model) where the transformation was performed is indicated at the top of the plot. The reported values are averages over three simulations while error bars denote standard errors of the mean. An asterisk (*) indicates a difference that is statistically significant (p-value calculated from a Student’s t-test smaller than 0.05) while a plus (+) indicates a difference that is marginally statistically significant (p-value between 0.05 and 0.10). (b) The estimated ΔΔG of binding due to the conversion of an individual methionine residue to methionine sulfoxide (see “Materials and Methods” and Figure 2). A positive value indicates that oxidation of a specific methionine residue is thermodynamically unfavorable for binding. Error bars in (b) are derived from the er/ror values displayed in (a) using the error propagation formula: 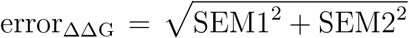, where SEM1 and SEM2 are the standard errors of the mean of ΔG_alch._^1^ and ΔG_alch._^2^, respectively.

**Figure 7:**
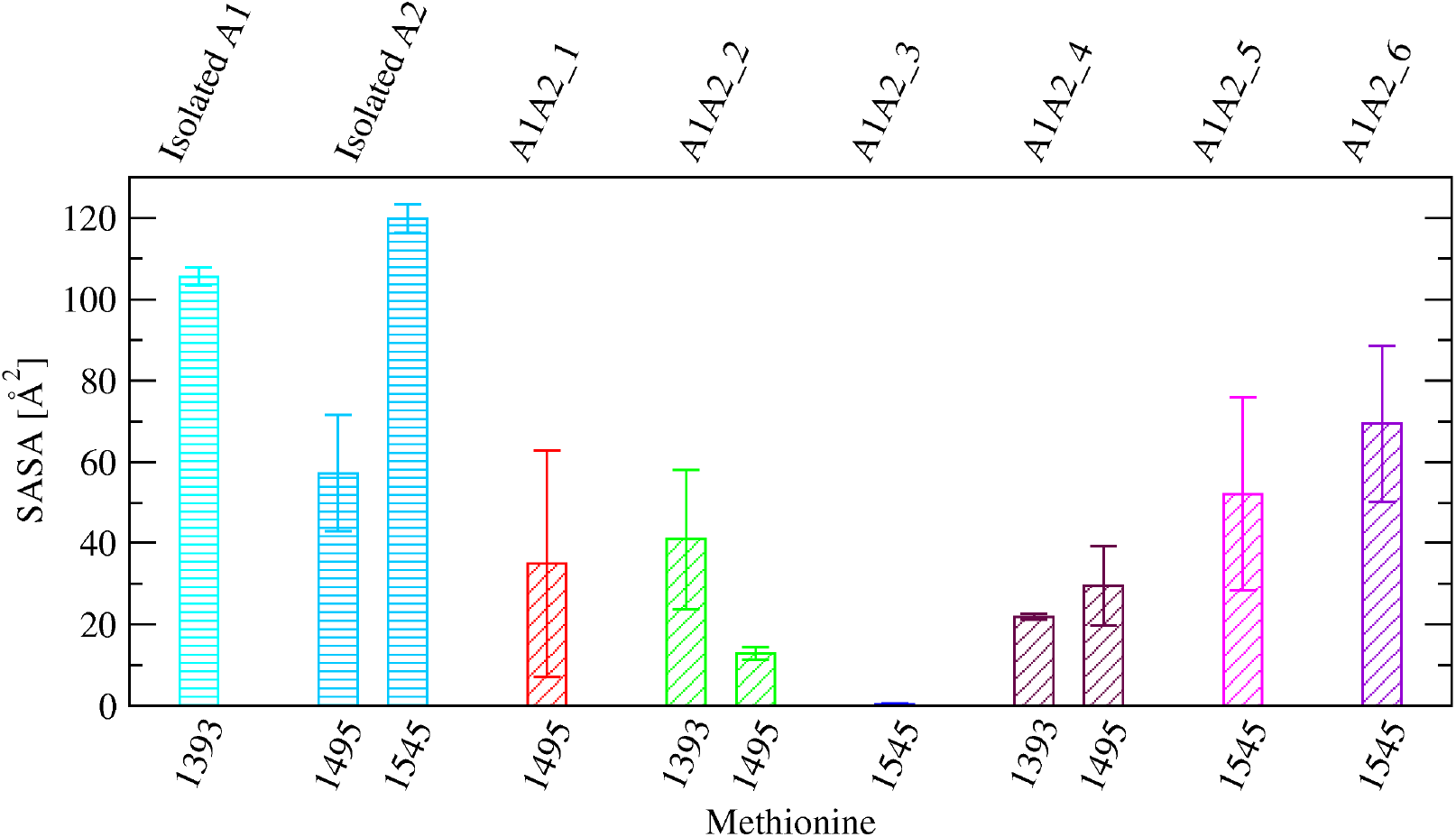
SASA of methionine residues located at the inter-domain interface. Comparison of SASA values of specific methionine side chains between the isolated A1 or A2 domain, respectively (bars with horizontal lines), and the A1A2 complex models where the specific methionine residue is located near the interface (bars with oblique lines). Reported are averages and standard errors of the mean over two simulations for the isolated A1 domain and the complex models, and three simulations for the isolated A2 domain, respectively.

## Discussion

The VWF protein is one of the key elements that bridge inflammation and thrombosis. Through the respiratory burst, neutrophils produce H_2_O_2_, which is converted by the action of MPO to HOCl. The latter has been shown to increase the oxidation of methionine residues in the A1, A2 and A3 domains of VWF especially under shear stress conditions.^4^ Flow chamber experiments performed in this study indicate that the presence of HOCl reduces the inhibitory function that the A2 and A3 domains normally have on the A1 domain, which is then more likely to bind to GpIb*α* on the surface of blood platelets. Computational models of the complex between the A1 and the A2 domains determined in a previous study^23^ provide evidence that the A2 domain is likely to obstruct the GpIb*α* binding site in the A1 domain. Furthermore, we found that in all models, at least one methionine residue in the A1 or A2 domain was located at the inter-domain interface (Figure 3). Calculations performed here based on FEP and the thermodynamic cycle estimated that oxidation of methionine residues located at the inter-domain interface is likely to destabilize the bound state of the A1 and A2 domains. The destabilizing effect of methionine oxidation was observed to be strongest when a methionine residue was entirely buried at the interface although it was still significant even when a methionine was partially buried (Figure 6b). In electron-microscopy images of dimeric VWF, the A1 domain was observed to adopt variable positions with respect to the A2 domain.^45^ Thus it is not excluded that multiple binding modes are possible between A1 and A2 and methionine oxidation reduces the number of accessible binding conformations. Taken together, these observations suggest that methionine oxidation reduces binding between the A1 and A2 domain facilitating the exposure of the GpIb*α* binding site in the A1 domain.

A previous study performed by us indicated that methionine oxidation also destabilizes the fold of the A2 domain and makes it more susceptible to tensile force.^21^ The fold of A2 differs from that of A1 and A3 in that it does not contain a disulfide bond linking the terminii. For this reason, tensile force due to shear forces present in rapidly flowing blood cause the A2 domain to unfold and expose a site that under normal non-oxidizing conditions is cleaved by ADAMTS13.^26^ However, the presence of HOCl has been shown to oxidize a key methionine residue in the cleavage site abrogating proteolysis through ADAMTS13.^5^ In summary, two mechanisms are likely to be present how methionine oxidation removes the inhibitory function of the A2 domain: facilitating unfolding of A2 and directly disrupting the binding interface with the A1 domain.

Both observations, destabilization of the A2 domain fold and reduced binding affinity between A1 and A2 due to methionine oxidation, have direct implications for the development of drug molecules aimed at preventing or reducing the risk of thrombosis. Currently available anti-thrombotic therapeutics carry the risk of haemorrhagic complications such as intracranial bleeding. For this reason, the development of a therapeutic that inhibits VWF selectively under conditions that lead to pathological thrombus formation would reduce the risk of thrombosis while still maintaining the normal haemostatic response. Knowledge of the atomistic details how methionine oxidation unmasks the GpIb*α* binding site in the A1 domain is necessary to identify sites that can be targeted selectively under oxidizing conditions. The results obtained here suggest for example the design of a molecule that binds to methionine residues located at the A1A2 inter-domain interface only after they are converted to methionine sulfoxide thus stabilizing the A1A2 complex and preventing binding of A1 to GpIb*α*.

In this manuscript, we propose that methionine oxidation shuts down an auto-inhibitory mechanism in the multi-domain VWF protein. Methionine oxidation is likely to play a similar role also in many other multi-domain protein systems. For example, methionine oxidation has been found to activate calcium/calmodulin-dependent protein kinase II in the absence of calcium^46^ and to activate I*κ*B*α*, an inhibitor of NF-*κ*B transcription factor.^47^ It is plausible that in these examples, methionine oxidation also removes an inhibitory mechanism that domains exert onto each other similarly as in VWF. In general, methionine oxidation regulates many other multi-domain proteins in the blood favoring a pro-thrombotic state such as ADAMTS13,^48^ thrombomodulin (TM),^49^ protein C^50^ and fibrinogen.^51, 52^ Thus, the methods used in this manuscript may be applied to other systems that are also regulated by methionine oxidation.

In conclusion, this study proposes an explanation for the observed increase in platelet-binding activity of VWF under oxidizing conditions^4^ based on methionine oxidation shutting down an auto-inhibitory mechanism. Furthermore, we investigated effects of methionine oxidation on a multi-domain protein using a combination of a dynamic flow assay and FEP to estimate changes in binding affinity between two protein domains. This method may be applied to other multi-domain protein systems exposed to shear stress where post-translational modifications such as methionine oxidation play a regulatory role.

## Acknowledgments

We would like to thank Nichole Tyler for performing preliminary flow chamber experiments, Wendy Thomas for help with the analysis and interpretation of the results, Olga Yakovenko for help with the experimental setup, and An-Yue Tu for kindly providing the DNA constructs. The simulations were performed on the Comet supercomputer at the San Diego Supercomputing Center thanks to a XSEDE allocation^53^ with grant number TG-MCB140143, which is made available through support from the National Science Foundation. This research was financially supported by a NIH Career Development Award K25HL118137, a University of Washington Royalty Research Fund Award A132953, and a Phase 1 Technology Commercialization Award from the Washington Research Foundation to GI.

