## Supplemental data for "Oxidation shuts down an auto-inhibitory mechanism of von Willebrand factor"

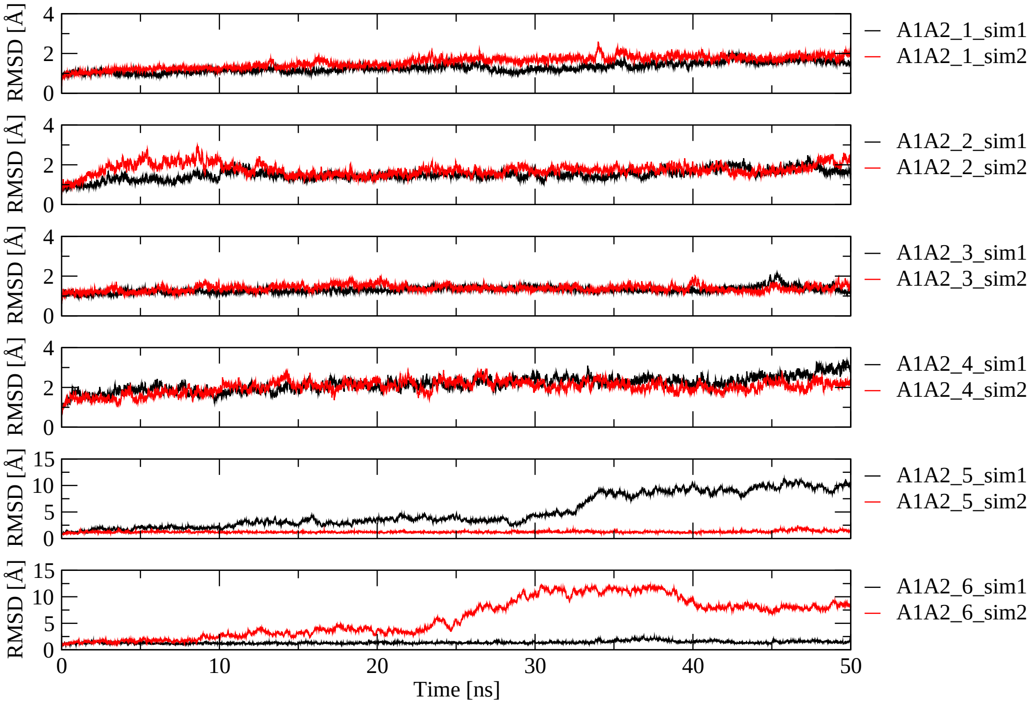


Figure S1: **Time series of the $C_\alpha$ RMSD from the initial conformation along the simulations with the A1A2 domains complex models.**

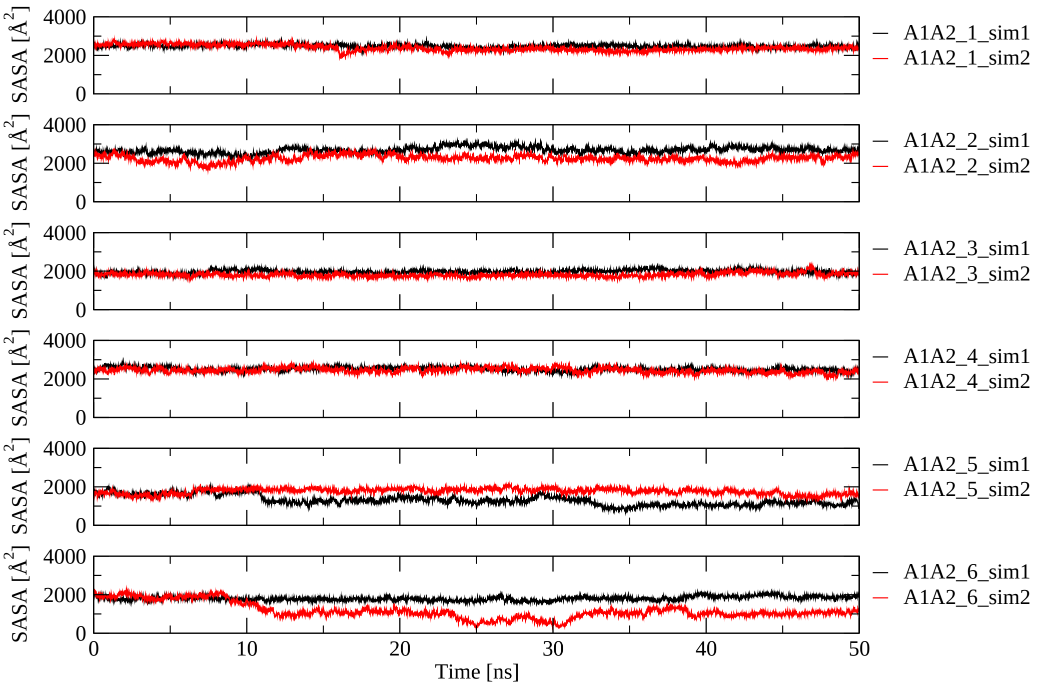


Figure S2: **Time series of the SASA buried at the inter-domain interface along the simulations with the A1A2 domains complex models.**
